# Astroglial Dysfunction, Demyelination and Nodular inflammation in Necrotizing Meningoencephalitis

**DOI:** 10.1101/2024.11.11.623107

**Authors:** Sara I Hernandez, Charles A Assenmacher, Molly E Church, Jorge I Alvarez

## Abstract

Necrotizing Meningoencephalitis (NME), a form of Meningoencephalitis of Unknown Origin (MUO), is a progressive neuroinflammatory disease that primarily affects young, small-breed dogs. Due to limited understanding of its pathophysiology, early detection and the development of targeted therapies remain challenging. Definitive ante-mortem diagnosis is often unfeasible, and dogs with NME are frequently grouped under the broader MUO category. Our long-term objective is to identify distinct disease mechanisms within each MUO subtype to improve diagnostic accuracy, therapeutic approaches, and prognostic outcomes. To establish unique inflammatory patterns as they relate to neuropathologic changes in NME, we studied we studied the degree of immune cell infiltration, astrogliosis, demyelination, and microglial activation, comparing these factors with granulomatous meningoencephalomyelitis (GME), a closely related MUO subtype. We found that in the leptomeninges, NME is characterized by mild immune cell infiltration, in contrast to the prominent, B cell-rich aggregates seen in GME. In the neuroparenchyma, both diseases exhibit a comparable degree of lymphocyte infiltration; however, demyelination is more pronounced in NME, particularly within the subcortical white matter. Notably, areas of the brain affected by NME display a reduction in astrogliosis, which is associated with a marked decrease in the expression of the water channel protein aquaporin-4 (AQP4), a reduction not observed in GME. Additionally, we found that AQP4 expression levels correlate with the extent of microglial and macrophage activation. These findings suggest that astrocyte dysfunction in regions of microglial inflammation is a driver of NME and with adaptive immune responses likely playing a supportive role.

## Introduction

Meningoencephalitis of Unknown Origin (MUO) refers to a group idiopathic neuroinflammatory diseases in canines. They include Granulomatous Meningoencephalomyelitis (GME), Necrotizing Meningoencephalitis (NME), and Necrotizing Leukoencephalitis (NLE). Clinical diagnosis of MUO is made by ruling out infectious, neoplastic, or other neurologic diseases affecting the Central Nervous System (CNS) ^1^. The treatment for MUOs typically involves immunosuppressive drugs, such as corticosteroids, often combined with chemotherapy, along with supportive care to manage symptoms ^1^. While, signalment, MRI findings, and clinical signs are associated with specific MUO subtypes, definitive diagnosis of each MUO subtype is only made post-mortem by histopathological confirmation of distinct patterns of change in the brain ^2^. In GME, inflammation has classically been described as occurring in the white matter of the cerebrum, optic chiasm, midbrain, brainstem, and spinal cord, with a predilection toward caudal portions of the neuroaxis. Leptomeningeal and neuroparenchymal inflammation is characterized by the presence of aggregates of lymphocytes, plasma cells, and epithelioid macrophages extending from around blood vessels ^1&3^. Perivascular infiltration in GME is typically more pronounced compared to other MUOs^1,3&4^. Our group recently demonstrated that in GME the extent of B cell aggregation in leptomeningeal infiltrates correlates with the severity of sub-meningeal demyelination ^5^. NME mainly affects young, small breed dogs such as Pugs, Shih Tzus, Yorkshire Terriers, and Chihuahuas^1&6^. Genetic predispositions have been described in Pugs, Chihuahuas, Maltese, and other small breed dogs ^7,8&9^. Lesions are found more rostrally in the neuroaxis, mainly in the grey matter of the cerebral cortex and sometimes extend to the subcortical white matter ^1,4&6^. Leptomeningeal, neuroparenchymal, and perivascular inflammation is characterized by lymphocytes and macrophages, with some plasma cells documented. The characteristic histologic changes are typified by regions of parenchymal necrosis and rarefaction with associated activated macrophages. Moderate to large numbers of astrocytes are often observed at the periphery of these lesions ^3&10^. In NLE, Yorkshire Terriers and French Bulldogs are overrepresented though it occurs in other small breed dogs ^1^. Necrotizing lesions of NLE are localized more rostrally in the neuroaxis, like NME, but are found mainly in the subcortical white matter with associated activated macrophages. In contrast to GME and NME, there is minimal leptomeningeal and neuroparenchymal lymphoplasmacytic inflammation ^1&6^.

Dogs diagnosed with MUO have a guarded prognosis. Moreover, those with NME typically have a rapid decline after even with treatment ^11^. Our understanding of NME immunopathogenesis is limited, though studies have identified a mutation in the canine major histocompatibility complex, also known as dog leukocyte antigen (DLA), in Pug, Chihuahua, and Maltese dogs that predisposes to NME ^9,12&13^. The DLA class I and class II antigen-presenting molecules play a key role in presenting non-self antigens to mount immune responses as well as regulate reactivity to self, a process essential for maintaining tolerance ^9,12&13^. Interestingly, there is evidence supporting an autoimmune component in NME, as autoantibodies against Glial Fibrillary Acidic Protein (GFAP), an intermediate filament protein expressed by astrocytes and upregulated during inflammation or injury, have been detected in the CNS of affected dogs ^14 & 15^. Given the limited understanding of the pathophysiology of MUO, particularly NME, this study aimed at elucidating the role of immune-mediated components in driving the neuropathology of NME.

## Methods

### NME and GME Cases

The archives of the Penn Vet Diagnostic Labs at the Ryan Veterinary Hospital of the School of Veterinary Medicine were searched for NME and GME cases diagnosed post mortem by ACVP board certified pathologists based on signalment, clinical signs, and histologic findings ^5^. Formalin-fixed and paraffin embedded (FFPE) blocks from NME (*n* = 11) and GME (*n* = 11) cases were utilized for this study (Table 1). FFPE blocks sampled from the cerebrum from five neurologically normal dogs were utilized as controls (Table 1).

**Table 1.** Clinical features of cases studied.

|  | **Breed** | **Age** | **Sex** | **Treatment** | **Clinical Presentation** |
| --- | --- | --- | --- | --- | --- |
| NME | Pug | 11 | MC | Keppra & Phenobarbital | Seizures |
| NME | Chihuahua | 3 | MC | Keppra, Prednisone & Cytarabine | Focal & tonic-clonic seizures, absent menace response, decreased proprioception & behavioral changes |
| NME | Mixed | 5 | MC | Prednisolone | Vestibular signs, decreased proprioception, absent pupillary light reflex & absent menace response |
| NME | Mixed | 8 | MC | Midazolam & Phenobarbital | Cluster seizures, absent proprioception, non-ambulatory & absent menace response |
| NME | Pug | 1 | FS | Phenobarbital & Dexamethasone | Generalized & focal seizures, absent menace response & decreased proprioception |
| NME | Yorkshire Terrier | 10 | FS | Keppra, Prednisone, Midazolam, Phenobarbital & Cyclosporin | Seizures, non-ambulatory tetraparesis, absent proprioception & absent menace response |
| NME | Mixed | 6 | MC | Keppra, Phenobarbital, Diazepam, Mannitol, Midazolam & Dexamethasone | Grandmal seizures & absent menace response |
| NME | Pug | 3 | FI | NA | Seizures |
| NME | Yorkshire Terrier | 0.5 | MI | Keppra, Phenobarbital, and Midazolam | Grandmal seizure, vocalizations, ataxia, circling, & head pressing |
| NME | Pug | 4 | FS | Keppra, Prednisone, Dexamethasone & Cyclosporin | Dull mentation, stargazing & lethargy |
| NME | Maltese | 2 | MC | NA | NA |
| GME | French Bulldog | 0.8 | MI | Prednisone, Rimadyl & Tramadol | Decreased proprioception & hind limb paresis |
| GME | Maltese | 6 | MC | Phenobarbital & Levetiracetam | Non-ambulatory, absent menace response & decreased proprioception |
| GME | Dachshund | 5 | FS | Cytarabine, Mannitol, Dexamethasone & Rimadyl | Head tilt, absent menace response & absent postural reactions |
| GME | Mixed | 8 | FS | Cytarabine, Mannitol & Dexamethasone | Vestibular ataxia, head tilt, cervical pain & circling |
| GME | Portuguese Water Dog | 10 | FS | NA | Circling, seizures & obtundation |
| GME | Labrador Retriever | 4 | MC | NA | Circling & vestibular ataxia |
| GME | Amer Pitbull Terrier | 7 | MC | NA | NA |
| GME | Shih Tzu | 5 | FS | Meloxicam & Prednisone | Ataxia |
| GME | Eurasier | 3 | MC | Phenobarbital & Diazepam | Epilepsy |
| GME | Amer Pitbull Terrier | 8 | FS | Prednisone, Dexamethasone & Mannitol | Head tilt, ataxia & absent menace response |
| GME | Mixed | 10 | FS | Prednisone | Vision loss & circling |
| Ctl | Labrador Retriever | 10 | MC | Dexamethasone, Methadone, Sucralfate & Maropitant | Gastric hemorrhage |
| Ctl | Schnauzer | 14 | MC | NA | Hepatocellular carcinoma & gallbladder mucocele |
| Ctl | American Bulldog | 3 | FI | Methadone, Maropitant & Ondasteron | Septic endometritis |
| Ctl | Staffordshire Terrier | 11 | FI | NA | Septic pyometria |
| Ctl | Labrador Retriever | 11 | MI | Maropitant, Butorphanol, Enrofloxacin, Acetylcysteine & Ampicillin | Fibrotic pleuropneumonia |

FS, female spayed; FI, female intact; MC, male castrated; MI, male intact; NA, not available; age in years.

### Tissue Processing and Immunostainings

Tissue samples were fixed for at least 24 h and FFPE blocks were sectioned 5-μm thick, mounted on glass slides, and stained with H&E. Tissue sections were processed for immunohistochemistry (IHC) as previously described ^5^. In brief, cortical areas of seven NME, eleven GME, and five control dogs were evaluated with antibodies (Abs) against CD3 (T cells) (M7254, Ms mAb; Agilent Dako), CD79b (B cells) (96024, Rb mAb; Cell Signaling Technology), Iba1 (macrophages/microglia) (019-19741, Rb pAb; WAKO), myelin-oligodendrocyte glycoprotein (MOG) (myelin) (96457, Rb mAb; Cell Signaling Technology), glial fibrillary acidic protein (GFAP) (GA52461-2, Rb pAb; Agilent Dako), and aquaporin-4 (AQP4) (AB3594, Rb pAb; Sigma Millipore). All the stains were performed using a LEICA Bond RXm automated IHC stain according to the manufacturer’s recommendations. Isotype control Abs were used as negative controls. Positive controls were normal canine brain tissue and lymph node. All NME, GME, and control IHC slides stained immunohistochemically were scanned at 20x using a Leica Aperio AT2 slide scanner (Leica Biosystems, Buffalo Grove, IL) and image acquisition was performed with ImageScope (Leica Biosystems, Buffalo Grove, IL). For the quantification of the MOG, GFAP, and AQP4 staining to assess demyelination, astrocyte activation, and AQP4 expression respectively. Image analysis was performed using a positive pixel count algorithm using ImageScope software. A single algorithm was developed using appropriate thresholds for the staining intensity and was applied to specific annotated regions on each slide.

### Neuropathological Analysis

Fulminant NME lesions are characterized by large areas of necrosis and neuroparenchymal loss mainly within the cerebral hemispheres. In contrast, GME is composed of single or multiple foci of inflammation mainly in caudal regions of the brain, including the spinal cord, though cerebral lesions can also be seen. To address the pathological differences between both diseases, our analysis was focused to areas of immune cell infiltration lacking necrosis and neuroparenchymal loss to reflect the potential early events leading to fulminant pathology in NME. Equivalent areas were also studied in GME and control cases. Leptomeningeal inflammatory infiltrates were scored using a four-tiered scale (absent= no inflammatory cells present; mild =fewer than 20 diffusely distributed inflammatory cells or leptomeningeal perivascular cuffing up to 3 layers; moderate = 20-50 diffusely distributed inflammatory cells or leptomeningeal perivascular cuffing three to seven layers; marked >50 diffusely distributed inflammatory cells or leptomeningeal perivascular cuffing greater than seven layers). Parenchymal cortical inflammation was scored using a similar scale: 0 =absent, 1 = mild (vessels with one cuff), 2 =moderate (many vessels with two cuffs), 3= marked (scattered or many vessels with >3cuffs). Evaluation of brain slides stained for for B cells (CD79b) and T cells (CD3) were based on previous methods ^5^. Glial nodules were scored using a four-tired scale (absent= 0 nodules; mild= 1 to 2 nodules; moderate= 3 to 4 nodules; severe= >5). To account for the variation in nodule size a three-tier scaled was added to the aforementioned score (0= 4 to 5 glial cells within the nodule; 1= 6 to 9 glial cells withing the nodules; 2= >10 glial cells withing the nodules.

### Statistical Analysis

Data were analyzed using GraphPad Prism v.10 software. Welch’ T test was used for comparisons between number of cell populations within leptomeningeal infiltrates. Brown-Forsythe and Welch ANOVA was used to determine the differential expression of MOG, GFAP, and AQP4 amongst control, GME, and NME dogs. Two tailed (95%) Pearson correlation analysis was used to determine associations between inflammation, lymphocyte infiltration, glial nodules, and algorithm outputs for the stains mentioned above.

## Results

### Mild leptomeningeal inflammation in NME

To investigate the underlying pathologic mechanisms of NME, we analyzed postmortem NME and GME cases with diagnoses confirmed post-mortem by a board-certified veterinary pathologist (MEC). As previously described, GME cases contained robust perivascular cuffing by lymphocytes ^5&16^. Histologically, perivascular cuffing was accompanied by adjacent aggregates as well as diffuse distribution of inflammatory cells within the leptomeninges (Figure 1A). In contrast, NME presented minimal perivascular immune cell aggregates within the leptomeninges and was characterized by a milder, diffuse infiltration of lymphocytes, which was most often associated with necrotic regions within the underlying brain neuroparenchyma (Figure 1B). Thus, NME and GME differ in the extent of leptomeningeal immune cell infiltration (Figure 1C).

**Figure 1.**
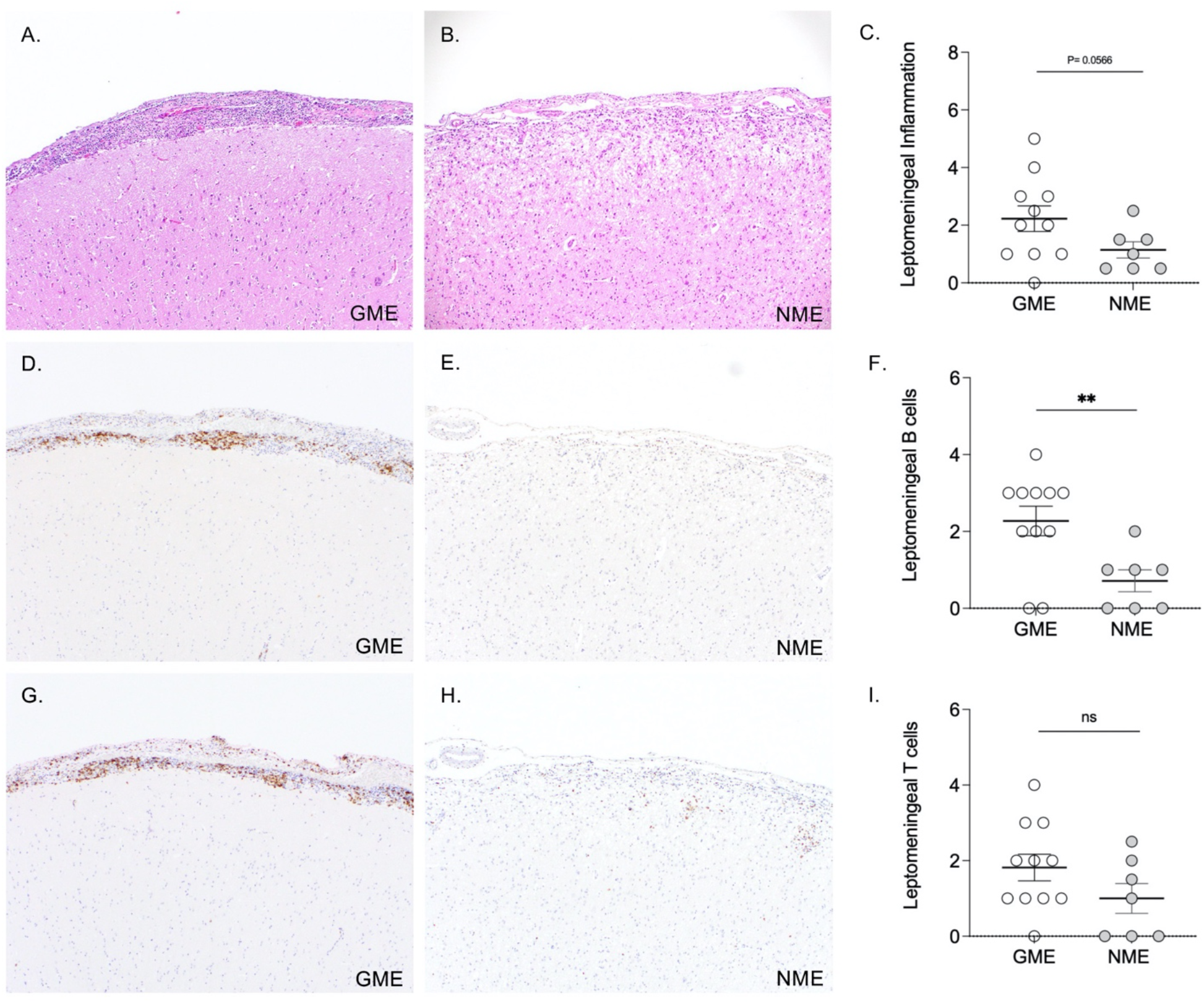
Leptomeningeal inflammation in NME and GME. (A & B) H&E stained slides of diffuse and perivascular inflammatory infiltrates at the leptomeninges. (C) Quantification of total leptomeningeal inflammation on H&E between GME and NME. (D & E) Leptomeningeal B infiltrates stained with anti-CD79b. (F) Quantification of total leptomeningeal B cells between GME and NME. (G & H) Leptomeningeal T cell infiltrates stained with anti-CD3. (I) Quantification of total leptomeningeal T cells between GME and NME. Tissues were counterstained with hematoxylin (blue) for nuclear staining. Error bars, mean ± SEM P ≤ 0.05* P≤0.01 ** by student t-test.

### Leptomeningeal B cells characterize GME from NME

Given that overall inflammation in the leptomeninges was less severe in NME than GME, we next determined whether the immune cell composition varied between both MUOs. As expected, the extent of B cell infiltration was significantly higher in GME (Figure 1D) while in NME limited number of B cells were distributed throughout the leptomeninges, mainly in a diffuse manner (Figures 1E-1F). We found similar levels of leptomeningeal T cell accumulation across both groups, though mildly higher numbers of T cells in GME compared to NME (Figures 1G-1I). These findings confirm a unique pathophysiological role for leptomeningeal B cells in GME while suggesting T cell infiltration may be similar in both pathologies ^5^.

### Comparable neuroparenchymal inflammation between NME and GME

The leptomeningeal inflammation in NME is mild and is without a predominant lymphocyte type, when compared to GME, where there tend to be more B cells present. When examining the inflammation within the neuroparenchyma in regions away from rarefaction and loss that are common in NME, there was no significant difference in the extent of cortical inflammation between the two groups (Figures 2A-2C). The neuroparenchymal lymphocyte composition of both NME and GME was largely similar (Figures 2D-2I). Therefore, while the composition of the inflammatory infiltrates observed in the neuroparenchyma do not differ between GME and NME, the leptomeningeal infiltration presents an enrichment of B cells in GME and limited lymphocyte aggregation in NME.

**Figure 2.**
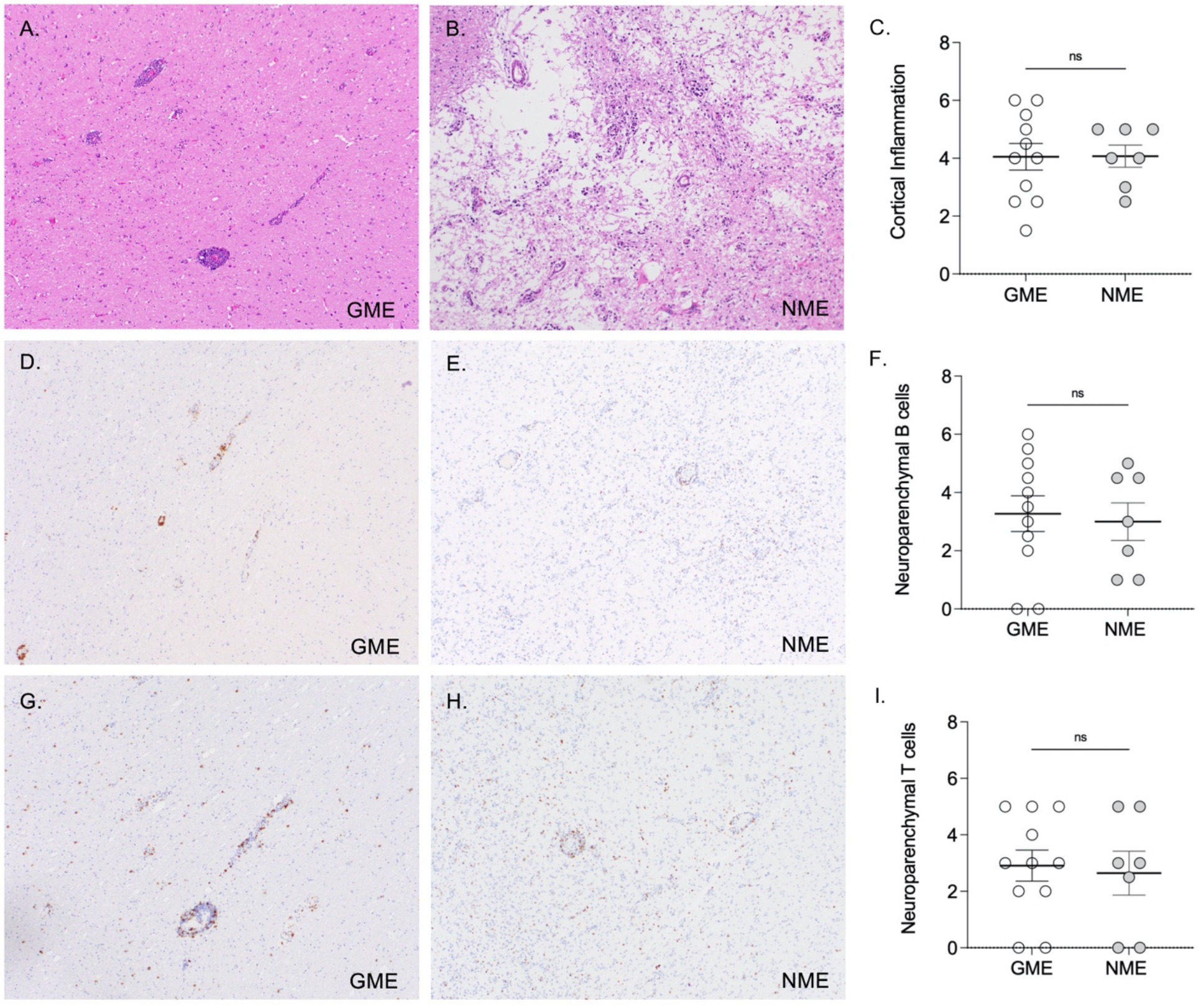
Neuroparenchymal inflammation in NME and GME. (A & B) H&E stained slides of diffuse and perivascular inflammatory infiltrates at the neuroparenchyma. (C) Quantification of total neuroparenchymal inflammation between GME and NME. (D & E) Neuroparenchymal B cells stained with anti-CD79b. (F) Quantification of neuroparenchymal B cells between GME and NME. (G & H) Neuroparenchymal T cell infiltrates stained with anti-CD3. (I) Quantification of total neuroparenchymal T cells between GME and NME. Error bars, mean ± SEM P ≤ 0.05* P≤0.01 ** by student t-test.

### Extensive subcortical white matter demyelination in NME

The presence of leptomeningeal B cells correlates to the extent of subpial demyelination in GME ^5^, and so we aimed to establish whether demyelination is also a component of NME. When compared to control groups, there was loss of MOG staining in the subpial, intracortical, leukocortical, and subcortical white matter layers of NME dogs (Figures 3A-3B). Specifically, demyelination was of the subpial grey matter and subcortical white matter in NME cases was greater than that present in GME cases (Figures 3C-3F). Sub-meningeal demyelination has been described as part of the pathophysiology of GME, however, distinct from GME, is the subcortical white matter demyelination in the cases of NME. Given the sparse lymphocyte infiltration observed in NME in the face of concurrent marked cortical and subcortical demyelination, we interrogated additional cellular components that may contribute to NME pathology.

**Figure 3.**
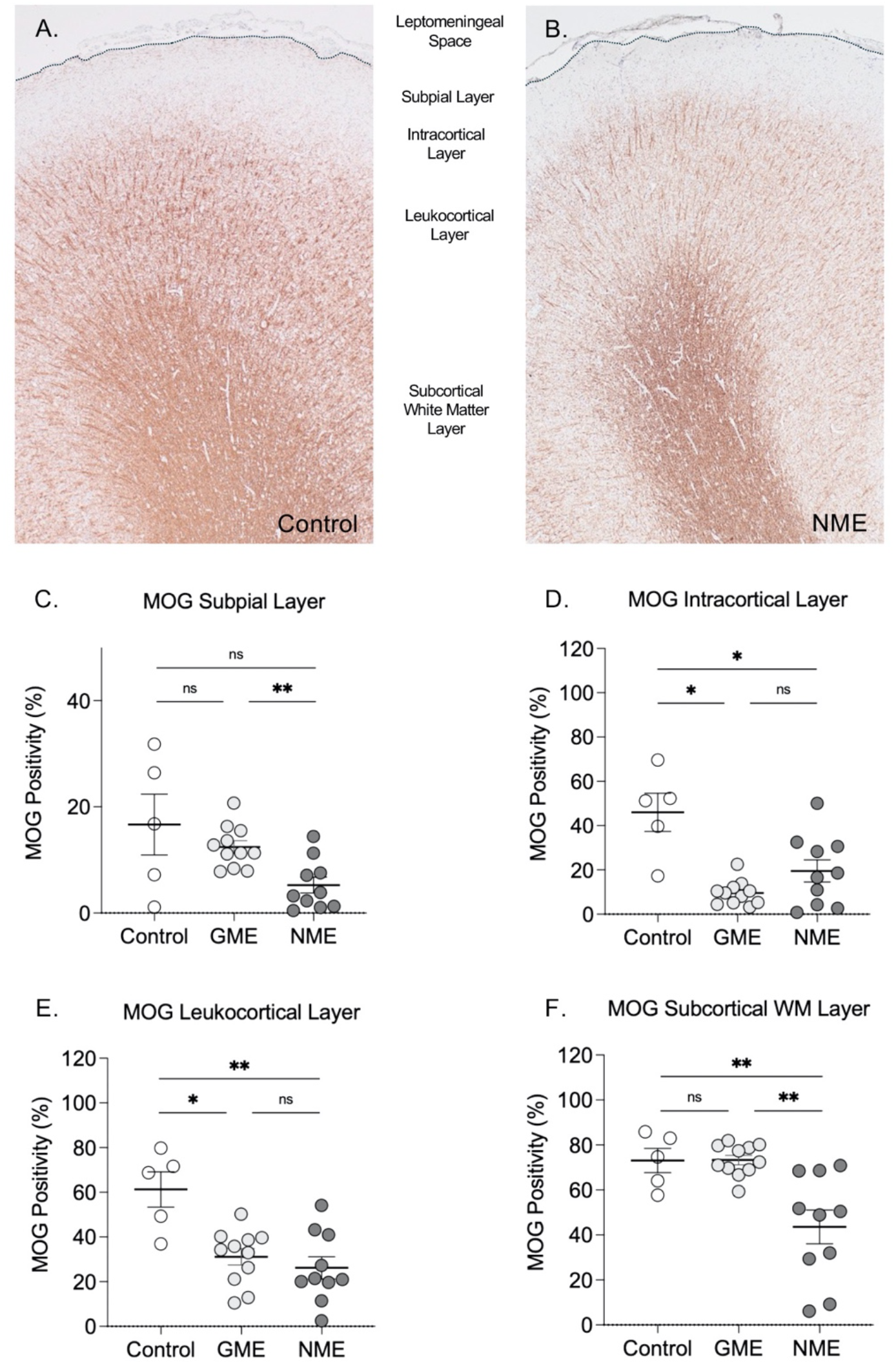
Demyelination in NME. (A & B) Representative images of control and NME dog brains stained with an anti-MOG antibody. The leptomeninges, subpial, intracortical, leukocortical, and subcortical white matter layers are indicated while the pia layer is labeled as a dotted lane. (C & F) Quantification of MOG expression between control (*n*=5), GME (*n*=11), and NME (*n*=10) across different brain layers. Error bars,mean ± SEM. *P≤0.05 ** P≤0.01 by one way ANOVA.

### Astroglial component in NME

Demyelinating and neuroinflammatory processes depend on astroglial activation to either propagate or subdue inflammation ^17^. We anticipated a higher extent of astrogliosis in NME as it has been suggested that antibodies against GFAP are present in CSF and serum of NME-affected dogs. These antibodies might potentially be targeting and subsequently activating astrocytes ^14 & 15^. Based on the considerable demyelination and the presence of inflammation, we suspected GFAP expression would be increased in both NME and GME at the cortical and subcortical level. We found that GFAP expression was significantly lower throughout the cortex and subcortical white matter in cases of NME when compared to control (Figures 4A-4F). In contrast, GFAP expression in GME was only significantly reduced at the intracortical and leukocortical layers (Figures 4D-4E). Throughout all layers GFAP levels trended downward in NME relative to GME (Figures 4C-4F). These findings indicate astrocytes might be a target of the inflammatory response seen in NME dogs.

**Figure 4.**
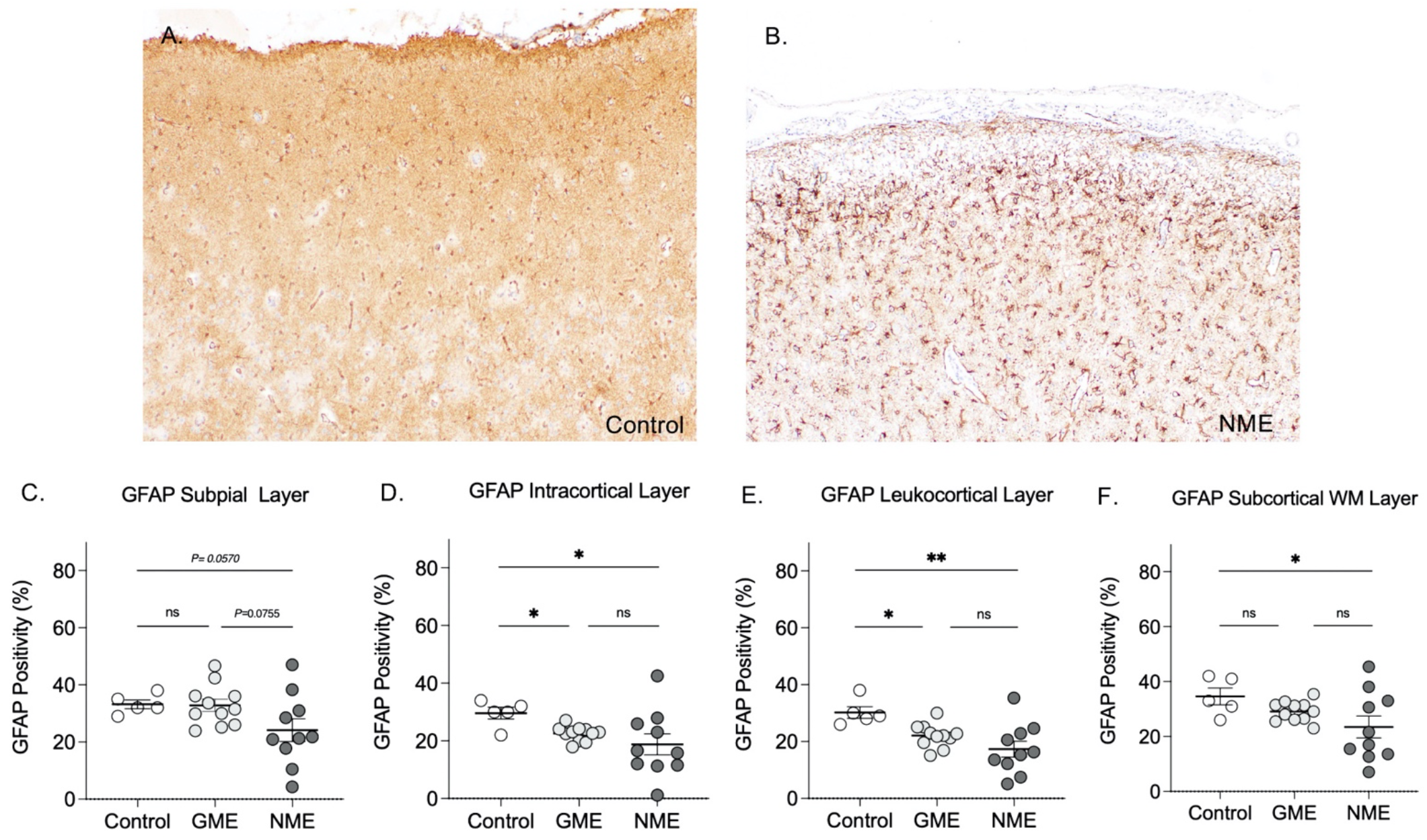
GFAP expression in NME. (A & B) Representative images of control and NME dog brains stained with an anti-GFAP antibody. Extent of GFAP expression at the subpial (C), intracortical (D), leukocortical (E), and subcortical white latter (F) in control (*n*=5), GME (*n*=11), and NME (*n*=10) brains. Error bars, mean ± SEM. *P≤0.05, ** P≤0.01 by one way ANOVA.

### Loss of AQP4 Distinguishes NME

As NME presents a reduction in GFAP expression, this led us to further examine the phenotype of astrocytes. We studied Aquaporin-4 (AQP4), a water channel protein expressed at astrocytic end-feet, involved in water homeostatic mechanisms, neuroexcitation, and astrocyte migration ^18^. The expression of AQP4 was reduced in cortex and subcortical white matter when compared to GME with significant decreases found in the subpial and leukocortical areas (Figures 5A-5C and 5E). The reduction of AQP4 expression was specific to NME (Figure 5C-5F). Therefore, these findings suggest that astrocytes are particularly affected and that AQP4 depletion is a distinct feature of NME pathology.

**Figure 5.**
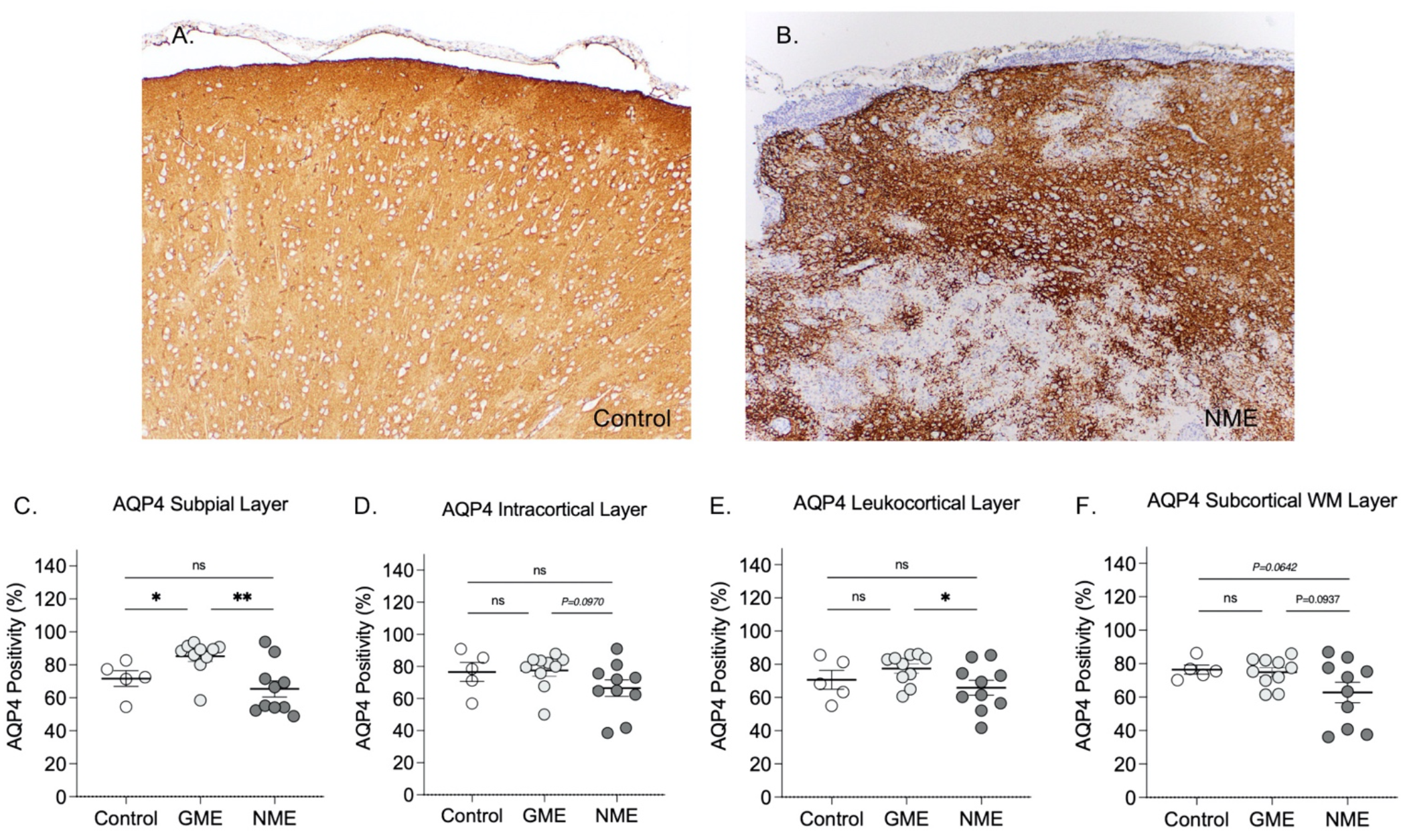
AQP4 expression in NME. (A & B) Representative images of control and NME dog brains stained with an anti-AQP4 antibody. Extent of AQP4 expression at the subpial (C), intracortical (D), leukocortical (E), and subcortical white latter (F) in control (*n*=5), GME (*n*=11), and NME (*n*=10) brains. Error bars, mean ± SEM. *P≤0.05 ** P≤0.01 by one way ANOVA.

### Loss of AQP4 correlates with decrease GFAP and increase demyelination in NME

The specific loss of AQP4 expression prompted us to determine if the demyelination observed in NME is associated with the extent of astrocyte damage across the cortex and within the subcortical white matter. In the cortex, we found no association between loss of MOG staining and the loss of GFAP staining (Figure 6A). Similarly, there was no correlation between the expression levels of GFAP and AQP4 (Figure 6B) nor between expression of MOG and AQP4 in cortical layers (Figure 6C). In contrast, within the subcortical white matter loss of GFAP expression was significantly correlated with a decrease in MOG expression (Figure 6D). Additionally, similar associations between AQP4-GFAP and AQP4-MOG expression (Figures 6E-6F) suggests that damage to both astrocytes and oligodendrocytes specifically within the subcortical white matter, may be related to a decrease in AQP4 expression as well.

**Figure 6.**
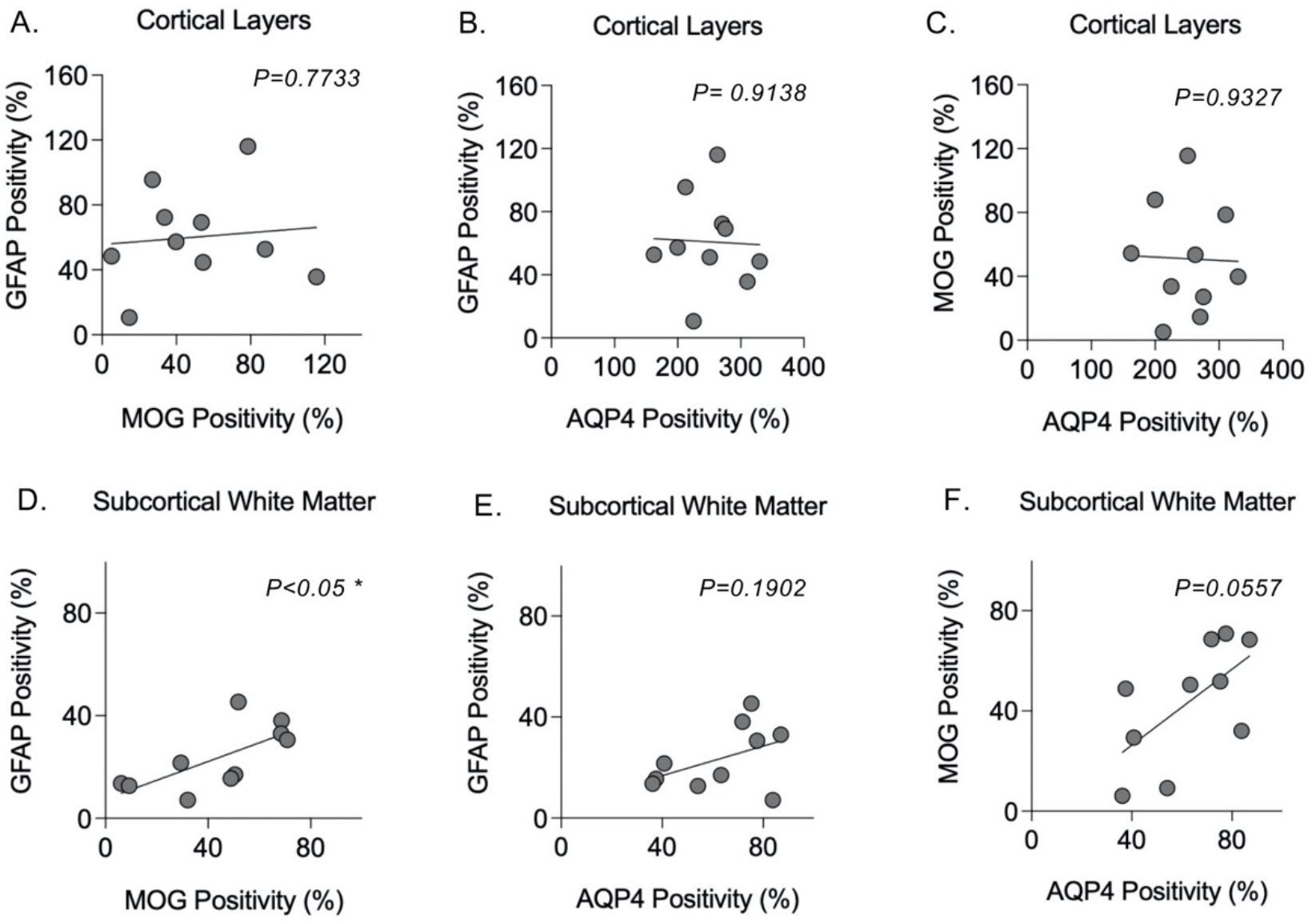
AQP4 loss in the white matter correlates with demyelination. We performed correlation analysis to determine the interdependence between astrocyte activation, demyelination, and AQP4 loss in the cortex (A-C) and subcortical white matter (D-F) of NME dogs (*n*=10). *P≤0.05

### Absence of AQP4 correlates to glial nodule and immune cell presence

We also examined if microglial activation correlated with loss of AQP4 observed in NME. We used an antibody against Iba1 to detect microglia, the resident innate immune cells of the CNS, and macrophages which typically migrate into the CNS from the periphery in response to inflammation. We hypothesized that the severity of microglia/macrophage activation would be associated with loss of both MOG and AQP4 within the neuroparenchyma. Glial nodules were observed in both the cortical and subcortical white matter, varying in the size according to the number of cells they contained. These nodules ranged from small clusters of four cells to larger structures with more than ten cells per nodule. We found no significant association between the presence of glial nodules and loss of MOG staining in the cortical layers (Figure 7A). However, within the subcortical white matter, expression of MOG trended towards being decreased with increased number of glial nodules. Expression of AQP4 was negatively correlated with the presence of glial nodules in the cortical layers and trended towards a correlation in the subcortical white matter layer (Figures 7C-7D). This suggests that a change in function of astrocytes in regions with nodular inflammation, away from homeostasis of the BBB and neuronal function and shifted towards inflammation.

**Figure 7.**
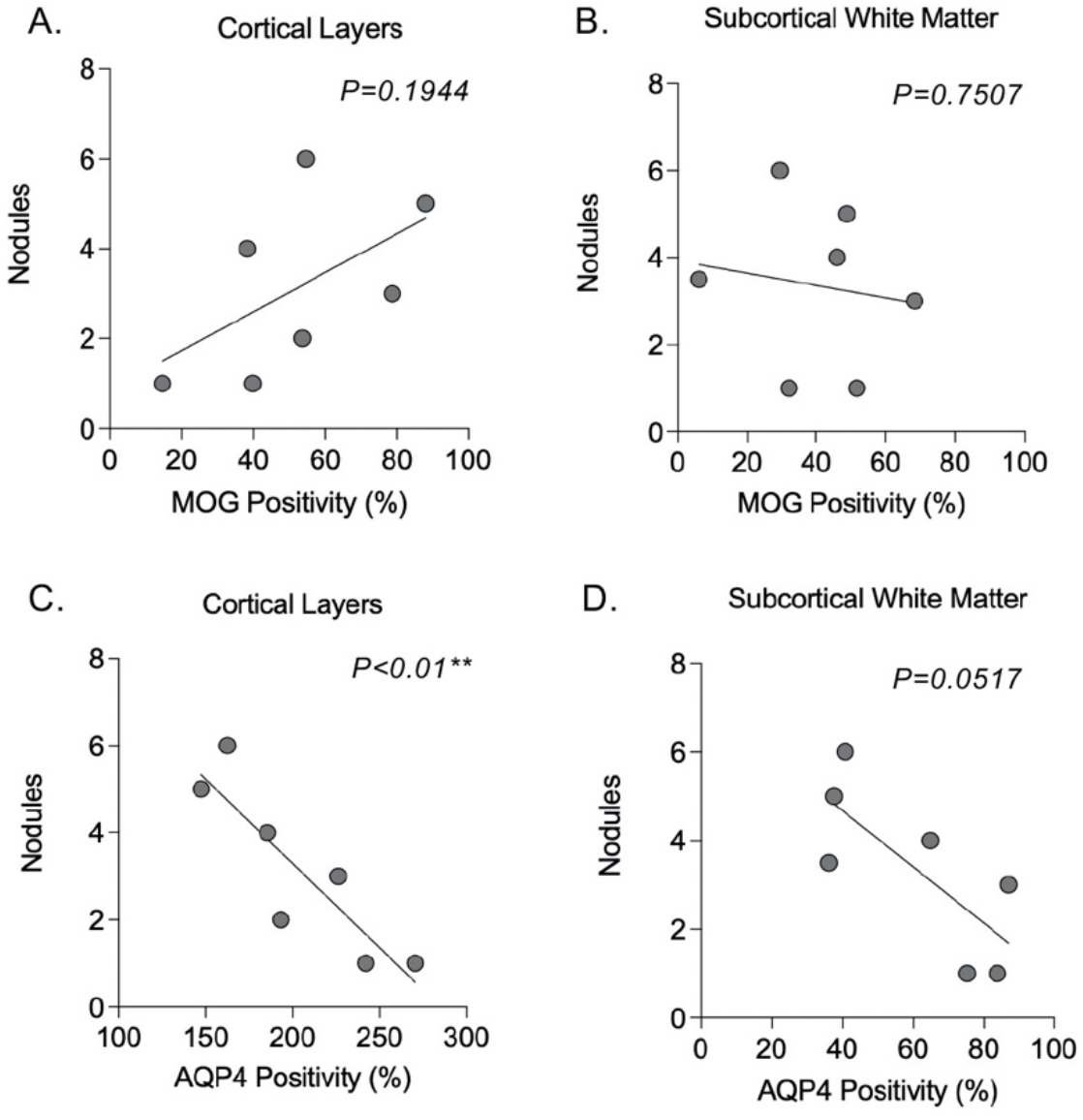
Microglial activation associates with the loss of AQP4 expression. We performed correlation analysis to determine the link between demyelination, and AQP4 with the extent of microglial activation (glial nodules formation) in the cortex (A-B) and subcortical white matter (C-D) of NME dogs (*n*=7). *P≤0.05, ** P≤0.01

This trend was also seen with severity of both B and T cell inflammation, where increased inflammation was correlated with decreased AQP4 expression in the cortex (Figures 8A-8B) but not in the subcortical white matter (Figures 8C-8D). Therefore, the presence of glial nodules was associated with loss of AQP4 in both layers, whereas the presence of lymphocytes correlated with AQP4 losses only at cortical layers.

**Figure 8.**
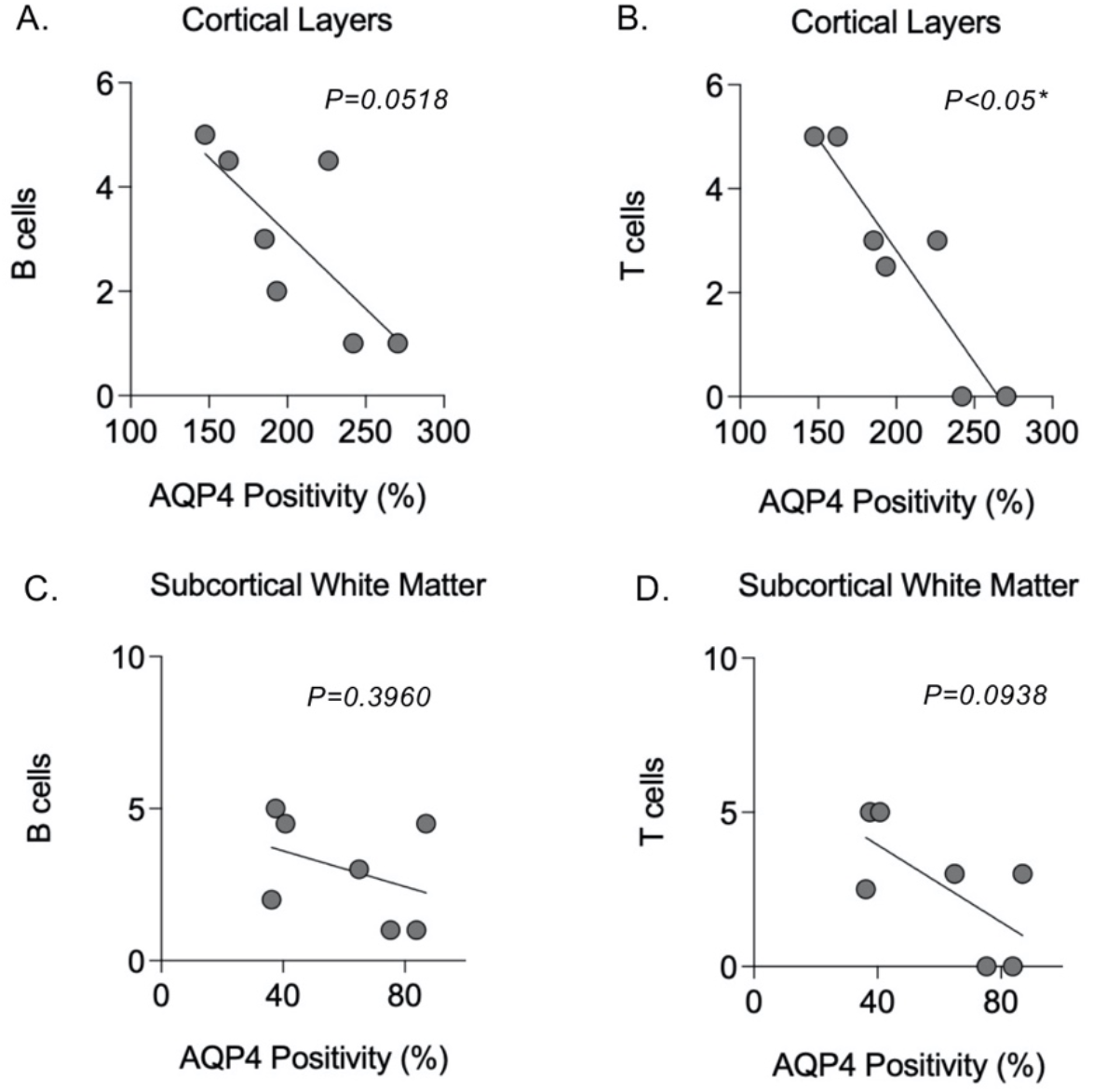
AQP4 loss correlates with the extent of lymphocyte infiltration in the cortex. We performed correlation analysis to determine the link between AQP4 and the extent of lymphocyte (B and T cells) infiltration in the cortex (A-B) and subcortical white matter (C-D) of NME dogs (*n*=7). *P≤0.05

## Discussion

While the histologic features of MUO subtypes have been well defined over the years, the underlying etiology of MUOs is largely unknown ^1,3,4,5,6&16^. Various hypotheses have been postulated regarding the mechanisms driving disease. Infectious etiologies have been suspected as causative, but none have been successfully detected in tissues ^19^. The most abundant body of evidence suggests NME to be either a primary autoimmune disease or a multifactorial immune-mediated disease. Multiple studies have found genetic mutations at the DLA Class II gene which potentially predisposes dogs to develop NME ^8,9,13,&20^. Moreover, there is also evidence of an autoantibody against GFAP identified in CSF and serum samples ^14,15, &21^, but it is still unknown whether these autoantibodies are directly causing the lesions or are secondarily produced from necrotic destruction exposing self-antigen, a mechanism known as epitope spreading ^22^.In this study, we provide evidence demonstrating variation in the immune-mediated mechanisms between NME and GME. Additionally, we identified unique features to the NME disease process such as demyelination and astroglial dysfunction.

Immune cell infiltrates comprising NME and GME lesions have been described in the past, though no studies have quantified the extent of immune cell infiltration in both the leptomeninges and neuroparenchyma. Similar to our findings here, T cells have been identified at the leptomeninges and neuroparenchyma of both NME and GME, with GME reported to have higher numbers of T cells, but with no reported statistical significance and in some cases with minimal presence in NME lesions ^4&6^. Notably, one group found no differences in B cell numbers in the neuroparenchyma between MUO subtypes, which is in accordance with our data^6^. In the GME cases examined, B cells were significantly more abundant than T cells in the leptomeninges and show a trend toward higher numbers in the neuroparenchyma. While in the NME cases, there were similar amounts of T cells in both compartments. The detection of both T and B cells in the leptomeninges and neuroparenchyma in NME suggests an ongoing inflammatory (including infectious, autoimmune, or immune mediated) disease process. Importantly, the presence of B cells in both NME and GME is indicative not only of chronicity, but signals the production of antibodies, which could then exacerbate a cyclical immune mediated process. Indeed, CSF from dogs with MUO has been demonstrated to contain CSF-specific oligoclonal bands (a metric of intrathecal immunoglobulin (IgG) ^10^.

As previously described by our group, leptomeningeal B cell aggregates are associated with submeningeal demyelination in GME cases ^5^. Although NME presented lesser leptomeningeal B cell infiltration, we interrogated whether demyelination was also present in association with immune cell aggregates. It is not widely agreed upon whether demyelination in NME represents a primary cause of the disease or a secondary response to loss of neuroparenchyma (and, thus, myelin) in cavitating lesions ^9&11^. Therefore, in this study we examined areas adjacent to necrotic lesions to determine the degree of demyelination. We did find extensive demyelination especially in subpial grey matter and subcortical white matter. Typical NME lesions are characterized by marked astrogliosis in and around necrotic regions ^1,3&6^. Astrocytes are the most abundant type of glial cells in the CNS, and they play a pivotal role in the maintenance of the neurovasculature as well as in nurturing neuronal function ^23 & 24^. During inflammation or injury, astrocytes become activated and increase in numbers as well as the expression of GFAP, a structural component mainly found at astrocytic processes ^25^. Our study contrasts the upregulation reported for GFAP during neuroinflammation and in combination with the focal loss of AQP4 suggest a potential immune targeting of astroglia unique to NME. If astrocytes are the direct or unintended target of antibodies, the resulting immune response could lead to astrocytic loss, which in turn would cause a reduction in the expression of GFAP and AQP4 ^26^. In this regard, the loss of AQP4 expression in NME correlated with the presence of microglial nodules, which form because of microglial activation needed to phagocytose neuroaxonal debris. Interestingly, the presence of microglial nodules does not correlate with demyelination in either cortical grey matter or subcortical white matter. In GME cases, demyelination was not associated with the presence of glial nodules but correlated with meningeal B cell infiltration ^5^. Therefore, microglial nodules might play distinct roles in each disease process. Microglia are involved in homeostatic functions (like synapse formation and neuronal survival) and immune mediated mechanism of the CNS such as phagocytosis of aged or degenerated myelin as well as other apoptotic cellular debris. Moreover, microglia are capable of promoting or downregulating neuroinflammatory responses ^27 & 28^.It is currently unclear whether microglial nodules are neuroprotective or neurodegenerative thus, further studies are required to elucidate the role of microglia in NME pathogenesis.

It is possible that the demyelination detected in NME is secondary to neuroinflammation. Reduced AQP4 expression also correlated with increases in B and T cell immune infiltration in the cortex. These findings mirror neuropathological findings seen in human neuroinflammatory disorders where the activation of B and T cells correlates with the presence of microglial nodules and ongoing CNS damage ^29&30^.Thus, primary astrocytic damage might be leading to a pro-inflammatory state, activation of microglia and perpetuation of inflammation. Such mechanism has also been postulated in humans with Autoimmune GFAP Astrocytopathy (AGA), a steroid responsive neuroinflammatory disease ^31^. Similar to NME patients, humans with AGA experience neck discomfort, seizures, personality/behavioral changes ^31^. Neuroparenchymal infiltrates in AGA consists of B and T cell with prominent microgliosis as well as loss of both GFAP and AQP4, and demyelination on histopathologic examination ^32^. Diagnosis of AGA is predicated on the presence of GFAP-specific IgG in CSF samples ^33^. Pre-mortem studies examining the specificity of IgG in CSF and serum of MUO dogs has the potential to identify a diagnostic biomarker and could aid in early diagnosis and targeted therapies for dogs suffering from NME.

## Acknowlegments

We want to thank the anatomic pathology residents and histology technicians of the School of Veterinary Medicine at the University of Pennsylvania for their help in procuring and processing tissues used in this study. We also want to thank the clinical veterinary neurologists for their contribution on the clinical diagnosis of these canine patients. This study has been supported by the The National Institute of Health (NIH) of the United States by the grant NCATS 5UL1TR001878-04 to the Institute of Translational Medicine and Therapeutics (ITMAT) of the University of Pennsylvania and JIA.

## Notes

### Competing Interest Statement

The authors have declared no competing interest.

